# The electro-MICA toolbox for integrating electrophysiology within multimodal imaging and connectomics workflows

**DOI:** 10.64898/2026.06.08.730888

**Authors:** Nicolás von Ellenrieder, Zhengchen Cai, Thaera Arafat, Laura Vavassori, Chifaou Abdallah, Jordan De Kraker, Raul Rodriguez-Cruces, Jessica Royer, Ella Sahlas, Paul Bautin, Raluca Pana, Olivier Aron, Birgit Frauscher, Boris C. Bernhardt

## Abstract

The integration of electrophysiological recordings with multimodal neuroimaging data holds great promise for advancing our understanding of brain function and neurological disorders. To facilitate this endeavour, we present electro-MICA, an open-access Python toolbox designed to project electrophysiological features from scalp and intracranial electroencephalography (EEG) onto cortical and hippocampal surfaces generated by validated multimodal imaging ecosystems. The toolbox comprises two pipelines: one for intracranial EEG (iEEG) recorded with stereo-EEG depth electrodes, and one for scalp EEG source localization. Both pipelines are grounded in numerical solutions to the electromagnetic equations governing electric activity in the brain, solved using the Boundary Element Method. A key methodological contribution is the use of a current density double layer model for neural generators, which avoids the mathematical singularities introduced by conventional dipole-based models when electrodes are near the cortical surface, a situation that can arise in iEEG. Electrode contacts are additionally modeled with non-zero length, improving physical realism. Scalp EEG source localization is performed using eLORETA on a subject-specific three-layer head model derived from the anatomical input. Validation against empirical gamma-band iEEG data from 32 subjects demonstrates that the distributed generator model outperforms both distance-based and dipole-based alternatives. An illustrative clinical example demonstrates the toolbox’s capacity to reveal associations between intracranial spike rates, cortical thickness, and anatomical connectivity in an epilepsy patient. Electro-MICA requires no parameter selection from the user, facilitating straightforward multimodal analyses in both research and clinical settings. The toolbox is available at github.com/MICA-MNI/electromica with extensive online documentation at electromica.readthedocs.io.

## 1. Introduction

The study of brain organization, function, and dysfunction has greatly benefitted from the availability and multimodal versatility of magnetic resonance imaging (MRI) (Alexander et al., 2007; Glasser et al., 2016; Halefoglu & Yousem, 2018; Larivière et al., 2019; Royer et al., 2022; Iutaka et al., 2023; Cabalo et al., 2025). These techniques provide structural and functional insights into the healthy human brain across different scales. They furthermore enable the precise delineation of lesions, malformations, and atrophic processes that underlie a broad spectrum of neurological disorders, such as neurodegenerative diseases, cerebrovascular pathologies, brain tumors, and epilepsy (REF). Multimodal neuroimaging, which integrates complementary information across these techniques, has further extended the sensitivity and specificity of structural assessments and is nowadays an essential step in advancing our understanding of both healthy brain function (Pagnotta et al., 2024) and pathologies (Calhoun & Sui, 2016). At the same time, advances in neuroinformatics and computational neuroimaging have enabled the registration and harmonization of diverse imaging contrasts into common reference spaces. Dedicated processing pipelines, such as FreeSurfer for cortical surface reconstruction and parcellation (Fischl, 2004) and HippUnfold for subfield-level hippocampal mapping (DeKraker et al., 2022), have made it possible to extract reproducible, biologically meaningful features from raw MRI data at scale.

Electrophysiological measurements such as electroencephalography (EEG) record the summated electrical activity of neuronal populations with millisecond-level temporal resolution, and constitute a foundation for the study of cognition (Dvorak et al., 2018), sleep (Berry et al., 2015), and evoked responses (Legatt, 2003) in healthy subjects, as well as electrophysiological abnormalities related to neurological pathologies such as Parkinson’s disease (Aljalal et al., 2022) or epilepsy (Noachtar & Rémi, 2009). The most common electrophysiological modality is scalp EEG, which measures changes of the electric potential on the scalp. Being non-invasive, low cost, and easy to standardize, it is appropriate for a broad range of applications. Intracranial EEG (iEEG), EEG recorded directly from the brain, is sometimes recorded in focal epilepsy patients who do not respond to pharmacological treatment. This iEEG recording is usually performed by implanting multi-contact EEG needle-like electrodes (stereo-EEG). It is clinically useful for determining the site of a possible surgical intervention, and at the same time provides an opportunity to record electric activity directly from the brain, useful for studying not only epilepsy (Duan et al., 2026) but also normal brain function (Frauscher et al., 2018). While scalp EEG and iEEG record the same underlying neurophysiological phenomena, they differ substantially in their spatial sampling. Scalp EEG captures broad, spatially blurred activity from large cortical regions (Von Ellenrieder et al., 2014), whereas iEEG has an extremely focal sensitivity (von Ellenrieder et al., 2021) and excellent signal-to-noise ratio (SNR). The temporal resolution of electrophysiological modalities and the spatial precision of anatomical imaging are naturally complementary, and their integration in multimodal datasets is an asset for the advancement of clinical neurology and the neuroscientific study of human brain function.

Here we present electro-MICA, a toolbox for the integration of electrophysiological information into a multimodal MRI processing and connectomics processing ecosystem. We specifically leverage micapipe (Cruces et al., 2022) which processes multimodal MRI data and maps these to cortical surfaces derived from FreeSurfer (Dale et al., 1999; Fischl, 2004)/FastSurfer (Henschel et al., 2020), as well as HippUnfold (DeKraker et al., 2022) for hippocampal segmentation and surface mapping. Electro-MICA is an open access, python toolbox with no dependencies other than python libraries. The toolbox projects electrophysiological features on the same cortical surfaces as the multimodal image processing results of micapipe, allowing for seamless integration. The electro-MICA toolbox consists of two pipelines, one for computing and presenting scalp EEG source imaging results on cortical/hippocampal surfaces, and another for projecting features associated with iEEG channels. In both cases our main objectives are ease of use and robustness, with no parameter selection necessary from the user. The in-house code computes numerical solutions of the equations governing electric phenomena in the brain, with a combination of unique models and proven algorithms.

## 2. Methods

We describe here the methods underlying each of the two pipelines of electro-MICA, namely scalp EEG source imaging and intracranial stereo-EEG. No parameters need to be selected by the user, making the pipelines extremely user friendly. This open access python toolbox is based on in-house code (github.com/MICA-MNI/electromica), with common packages for file handling and mathematical operations and it is documented at http://electromica.readthedocs.io. The Advanced Normalization Tools (Tustison et al., 2021) python package ANTsPy is required for coregistration of anatomical volumes associated with the EEG electrode positions to the *nativepro* space of micapipe. We obtained IRB approval for research in iEEG, high-density EEG, and MR imaging. Written informed consent was obtained from all participants included in the study.

### 2.1 Intracranial EEG pipeline

For iEEG data, the pipeline projects features associated with the iEEG channels onto the cortical and hippocampal surfaces produced by micapipe and HippUnfold to represent multimodal imaging outputs. This projection is informed by the physics governing the electromagnetic phenomena in the brain.

**Figure 1A** shows a diagram of the electroMICA_iEEG pipeline. The main input is a list of values for one or more features, associated with channels formed by the iEEG electrodes. This input is a simple table of tab separated values with the channel name and value for the features. The channels can be referential or bipolar. Additional inputs to the pipeline are the iEEG BIDS (Holdgraf et al., 2019) compliant electrode file and its associated volume (preferably a T1-weighted volume), the micapipe output (anatomical and surface folders), and optionally the HippUnfold surface folder output. The output of the pipeline is a map of each surface on the cortical and hippocampal surfaces, in GIFTI format (Harwell, John et al., 2011). An intermediate output of the pipeline, the sensitivity of each iEEG channel to electric activity localized around each vertex of the surfaces, is saved in the model folder. The rest of the pipeline output is populated with copies of the surfaces and anatomical images used. The electrodes and associated anatomical volume coregistered to *nativepro* space are also saved, as well as the corresponding transformation.

**Figure 1.**
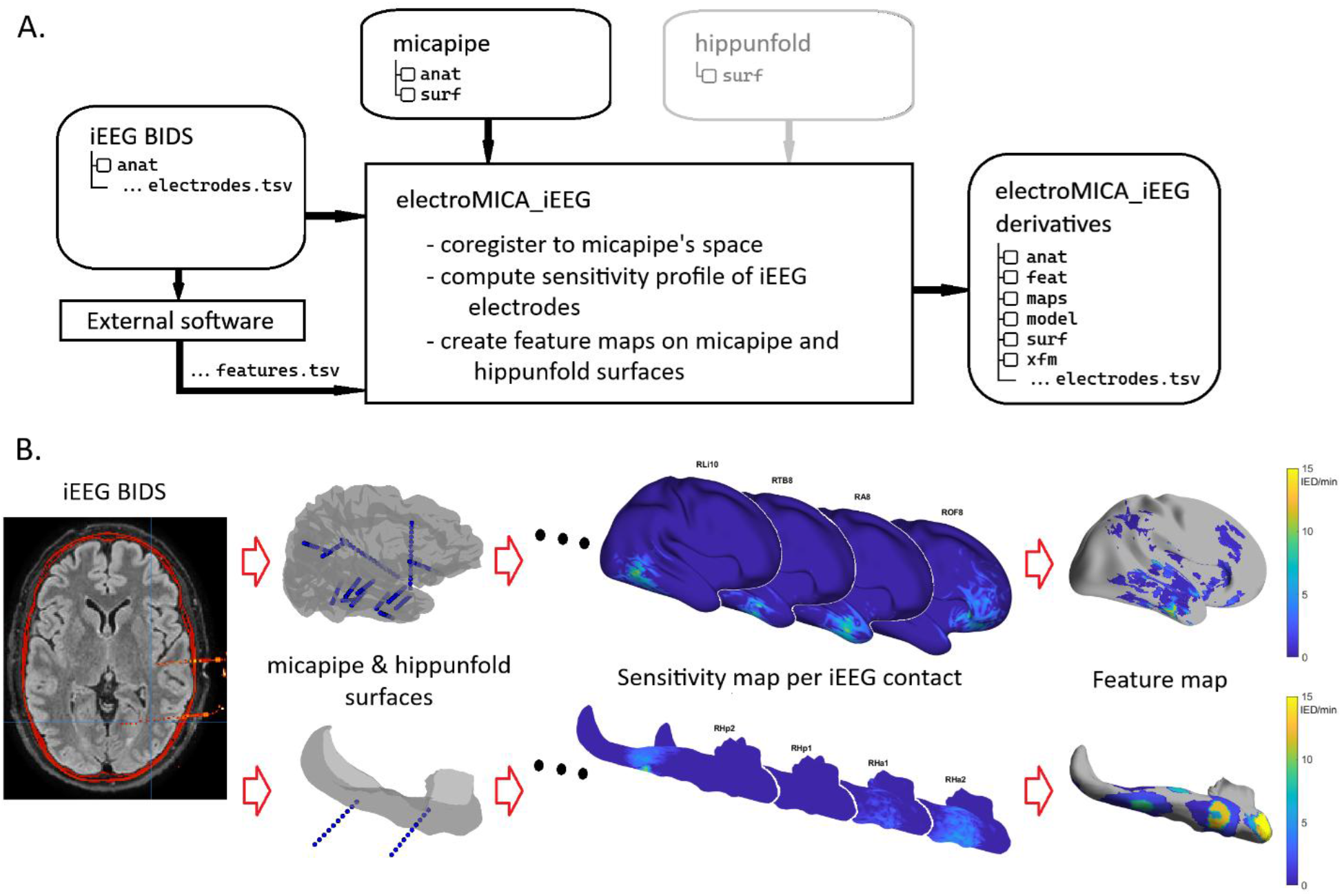
Schematic diagram of the intracranial EEG pipeline electroMICA_iEEG. **A** | Flow chart with inputs and outputs. Gray indicates optional features. **B** | Workflow representation: iEEG electrodes in BIDS format are coregistered with micapipe and hippunfold surfaces, then the sensitivity map of each iEEG contact is computed, and finally, contact sensitivity maps are combined and thresholded to obtain a representation of the input features of iEEG channels on the cortical surfaces.

**Figure 1B** depicts the main steps of the pipeline; an initial transformation of the electrode positions to micapipe’s *nativepro* space, the computation of the sensitivity of each iEEG channel to generators of electric activity on the cortical and hippocampal surfaces, and the construction of the maps associated with the input features. The first two steps need to be run only once for a given subject; additional maps can be quickly computed afterwards using the pre-computed models. To compute the sensitivity of the iEEG channels to neuronal generators of electric activity at each node of the cortical surface, the electromagnetism equations governing the electric phenomena in the brain are solved with the Boundary Element Method (BEM) (De Munck, 1992). The head is modeled as a single layer given by the inner-skull surface, obtained from the brain mask. The generators of electric activity are modeled as a current density double layer on the cortical surface, linearly interpolated between the nodes (Von Ellenrieder et al., 2009). Note that this model is different than usual models for distributed activity, which consist of a dipole at each node of the surface (Delorme & Makeig, 2004). While a model with dipoles is appropriate for computing the electric potential on the scalp, it involves mathematical singularities for the electric potential at the nodes, introducing large numerical instabilities when the electrodes are close to the cortical surface, as can be the case in iEEG. The more realistic and mathematical well-behaved model we adopt is unique to electro-MICA. Since the generator model has zero-thickness (*i.e*., a surface), we combine it with an electrode model of non-zero length of the metal contacts, as we expect it to improve the model. The metallic contacts of the electrodes are modelled as a line with the true length of the contact, not a single point. The electric potential at the electrode is computed as the average potential along this line (Von Ellenrieder et al., 2012), with 5^th^ order Gauss-Legendre quadrature (Golub & Welsch, 1969).

The sensitivity of iEEG decreases very rapidly in the brain with the distance to the contacts (von Ellenrieder et al., 2021). The sensitivity to generators far from the contacts is negligible compared to the measurement noise or masked by activity closer to the channel. Thus, two thresholds are applied for the constructions of the feature maps: a threshold common to all channels reflecting the effect of the noise (0.001 Vm/A), and a channel dependent relative threshold (0.05 of the maximum sensitivity of the channel). While these thresholds are arbitrary, they have been chosen based on extensive experience in iEEG data analysis. The sensitivity of each iEEG channel is computed based on the contact sensitivities (*i.e*., for a bipolar channel, it is the difference between the sensitivity of each contact), and then the thresholds are applied. Finally, each node of the surfaces is assigned the value of the feature from the channel with highest sensitivity at that node (piecewise constant output map) and a weighted average of the features with the weights given by the thresholded channel sensitivities (smoothed output map). Large portions of the cortex are typically far from all iEEG contacts and will not be observable with iEEG. The maps show no value (NaN) for the surface vertices in these regions.

### 2.2 Scalp EEG pipeline

In the case of scalp EEG, the pipeline performs source localization. The source space includes the cortical surfaces derived from micapipe and HippUnfold.

**Figure 2A** depicts the main input to electroMICA_scalp is a list of electric potential values that one or more features take for scalp channels in average reference format. This input is a simple table in tab separated values format. Additional inputs are the micapipe output (anatomical, surface, and transformation folders), and optionally the HippUnfold surface folder output. If the scalp channels are a subset of the 10/10 electrode position system (Nuwer, 2018), with the proper channel names, the position of the electrodes is assigned automatically. If the positions of the channels do not follow this system, an EEG BIDS (Pernet et al., 2019) compliant electrode file and its associated volume (preferably a T1-weighted volume) are also required.

**Figure 2.**
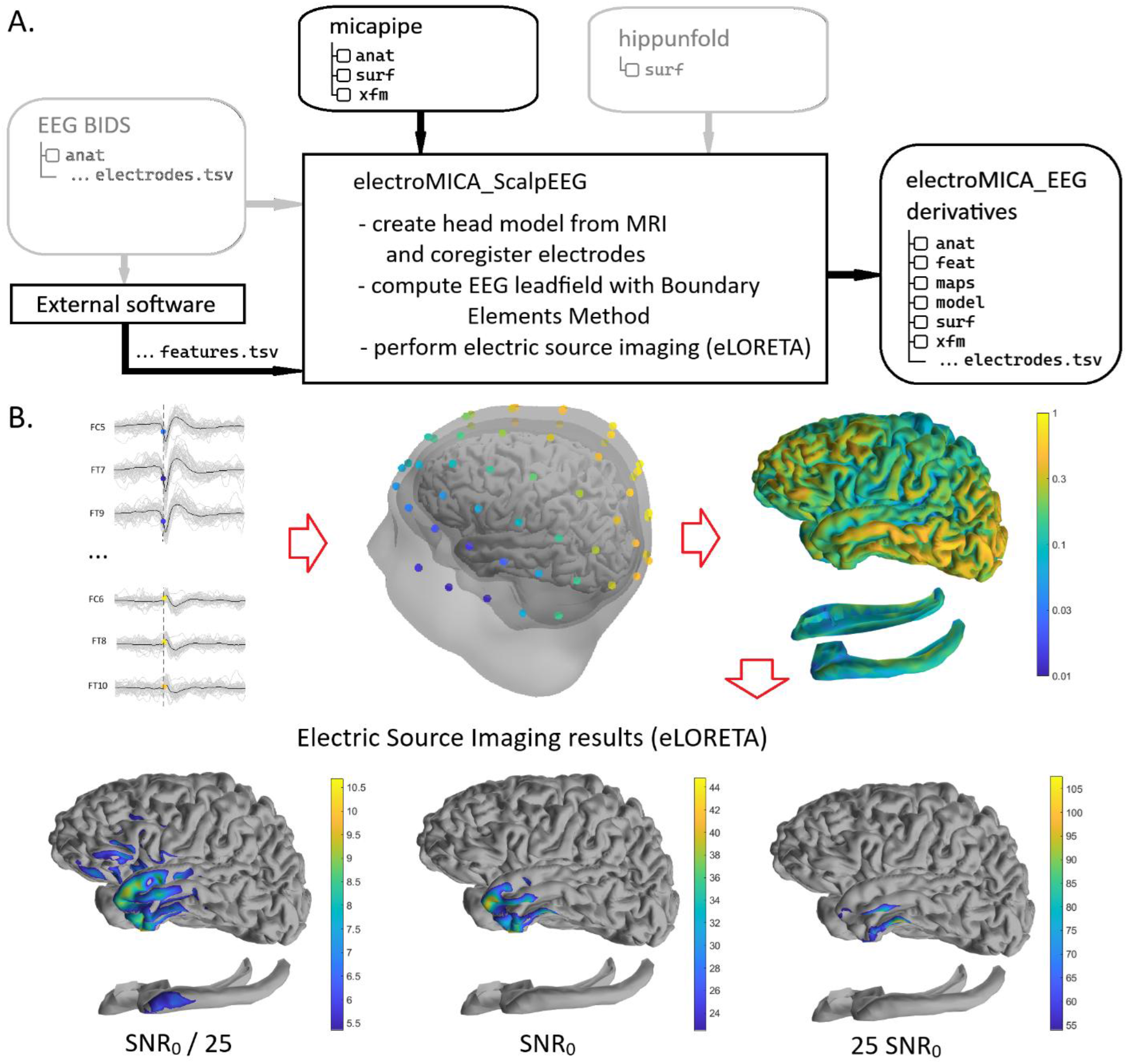
Schematic of the scalp EEG pipeline electroMICA_scalp. **A** | Flow chart detailing inputs and outputs. Gray color indicate optional inputs. **B** | Workflow of the pipeline. A set of values for scalp EEG channels in an average montage is the input; here, it is the electric potential amplitude of an average epileptic spike at the mid-point of the rising flank. The pipeline first builds a three-layer head model, then computes the sensitivity map of each channel, or leadfield. In the right center panel, the color on the cortex represents the total sensitivity, summed for all channels, of the EEG to a generator on that cortical location. Finally, the source imaging problem is solved using eLORETA, for different SNR ratios of the input electric potential. A low SNR (*left*) leads to more uncertain localization and lower source strength, while high SNR (*right*) leads to more certainty in the location and higher source strength.

**Figure 2B** shows the head model as a three-layer model (brain, skull, skin) built from the T1-weighted volume in *nativepro* space from micapipe and its brain mask, mainly through morphological processing. Anatomical landmarks in the pre-auricular points, nasion, and inion, as well as the location of 10-10 scalp electrodes are approximated from the transformed positions in MNI152 space, projected onto the scalp surface of the subject. If non-standard electrode locations are used, the anatomical volume associated with the BIDS compliant electrode file is coregistered to the micapipe anatomical volume. The generators of electric activity are modeled as a current density double layer on the cortical surfaces from micapipe, and optionally the hippocampal surfaces from HippUnfold, linearly interpolated between the nodes (*i.e*., the vertices of the tessellated surface mesh). The forward problem is solved using BEM, and the source localization using eLORETA (Pascual-Marqui et al., 2018), with a smoothed covariance prior. Five different solutions are computed for each feature, corresponding to very-low (SNR_0_/25), low (SNR_0_/5), medium (SNR_0_), high (5 SNR_0_), and very-high (25 SNR_0_) signal-to-noise values. The user can choose which solution corresponds to the data under analysis, from a single event almost completely masked by noise (very-low SNR), to an extremely clear average of a large number of events in a good quality recording (very-high SNR). No other parameters need to be selected by the user.

## 3. Results

The generator modeled as a current density on a surface is unique to this pipeline, and next, we explore its performance for iEEG modeling. In **Figure 3**, we provide examples of the sensitivity of a single intracerebral contact to generators on the cortical surface, shown on an inflated surface. Four models are compared for the generators. In the first case, the sensitivity is based simply on the Euclidean distance between the nodes of the surface and the center of the electrode contact, decaying with the inverse of the squared distance. This model does not take into account that the generated current is typically perpendicular to the cortical surface. The second generator model considers a mathematical dipole at each node of the cortical surface, with a normal orientation and a point model for the electrode, and it is the most widely used model. The third and fourth models are the ones with linearly varying current density double layer, with orientation normal to the cortical surface, the third with a point model for the intracranial electrode, and the fourth with a line model for it. In **Figure 3A**, we see an example in which the electrode is not very close to the surface, and all models produce similar sensitivity values. **Figure 3B** shows that the dipole-based model overestimates the maximum sensitivity and severely underestimates the extent of the 1% sensitivity region compared to the truly distributed models. **Figure 3C** corresponds to an example in which the center of the electrode contact is close to the cortical surface but far from the nodes, to illustrate the difference between the third and fourth models. The electrode modelled as a point leads, in this case, to a slightly higher maximum sensitivity and lower extent than a more realistic electrode model with non-zero electrode size. Finally, **Figure 3D** shows the correlation between the measured activity in the gamma band (30-100 Hz), and a relatively simple approximation of the expected signal based on the models, for all 3950 bipolar intracerebral channels of 32 subjects. We combine the data from all patients because the model has no subject-dependent parameters, *i.e*., the same physical equations govern the problem. The expected signal strength from the model is computed by assuming a normally distributed current density in each node of the cortical surface, with zero mean, constant variance, and independent between nodes. This is a very crude approximation, so a high correlation should not be expected even though for the gamma band the model is better than for lower frequencies, for which spatial correlation of the generators is larger (Gloor, 1985).

**Figure 3.**
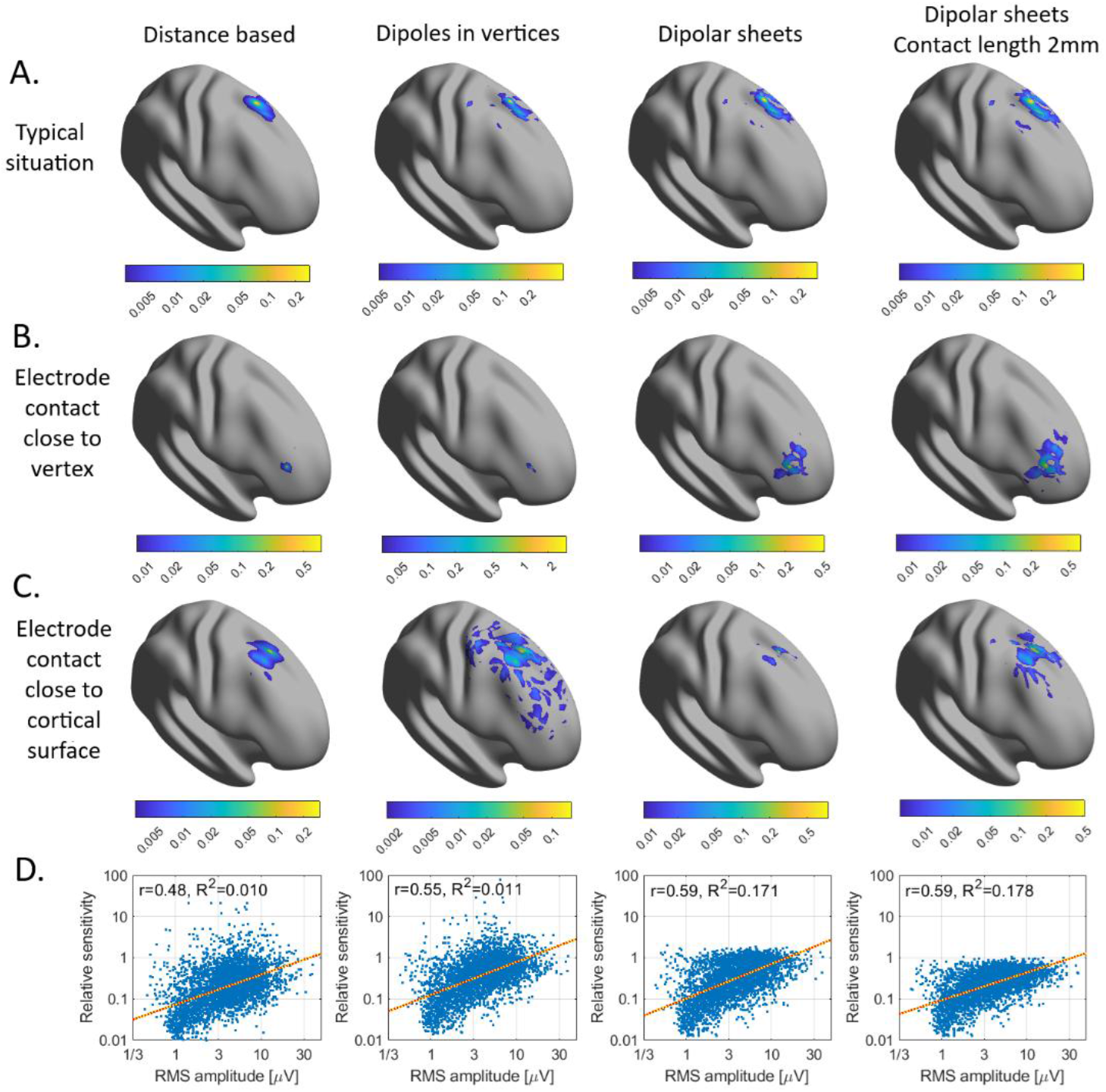
Examples showing the sensitivity maps obtained for intracranial EEG with different models for the source and electrode contacts. A model based only on the distance between cortex and electrode contacts (inverse of the squared-distance) is shown on the left, followed by a model with dipolar sources at the nodes of the cortical surfaces, and a model with a current density double layer on the cortical surface. The model on the right is also for a current density double layer as the source model, and a line model for the elctrode contacts, unlike the point model of the previous three cases. In all cases, the color indicates the sensitivity to a threshold of one hundredth of the maximum. **A** | For most source locations the models produce similar sensitivity maps. **B** | In cases when the electrode contact is close to a node, the first two models are numerically unstable due to inherent mathemtaical singularities, leading to extremely high maximum sensitivity and extremely small regions of non-negligible sensitivity. **C** | When the electrode contact is close to the cortical surfacebut not to a node, the physical dimensions of the contact play a role, since the whole line representing the contact is unlikely to be too close to the cortical surface. **D** | Correlation between the log-sensitivity of 3950 iEEG channels from 32 patients and the log-transformed measured EEG activity in the gamma band (30-100 Hz). The correlation and the proportion of explained variance is higher for the cases with a double layer current density source model.

While for all models the correlation is significant (p<=10^-5^, block permutation test, 10^5^ permutations), the options with the truly distributed generator model explain a higher proportion of the variance (R^2^ = 0.17 and 0.18) compared to the distance based and dipole-based models (R^2^ = 0.01). This supports our choice of models for the iEEG pipeline.

An example of how the pipeline allows for integration between information derived from imaging and from electrophysiology is shown in **Figure 4**. The data corresponds to a patient with temporal and posterior quadrant epilepsy, with 120 bipolar iEEG channels. The top row shows a comparison between the intracranial spiking rate and cortical thickness at corresponding sites. The image on the top right shows the rate of epileptic spikes of the iEEG channels projected on the cortical surface using electro-MICA. The spike rate was computed in a 10-minute interval during non-rapid-eye-movement (NREM) sleep, using an automatic spike detector (Janca et al., 2015). The top-center panel shows the correlation between the median of the cortical thickness z-score in the region from which each iEEG channel records and the log-spike rate for that channel. A significant negative Pearson correlation is observed (r=0.32; p=0.0094, block permutation test preserving the local spatial autocorrelation, 10^5^ permutations), as could be expected if there is some atrophy in mesiotemporal regions with high spike rates.

**Figure 4.**
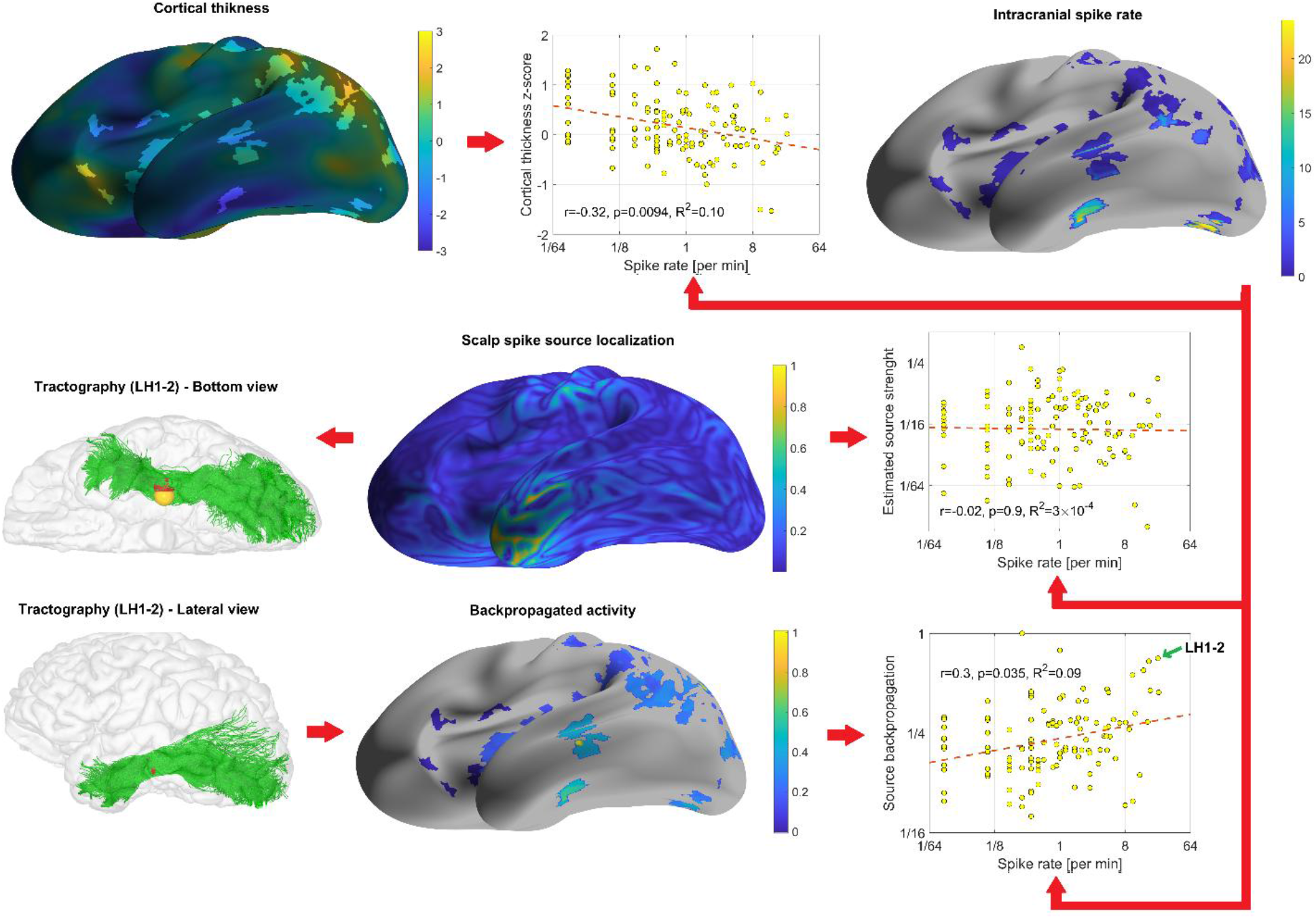
Examples of integration of electrophysiological and anatomical information, in a patient with temporal + posterior epilepsy. *Top*: correlation between the cortical thickness (z-score compared to healthy controls), at the locations from which the 120 iEEG channels record, and the epileptic spike rate for these channels. On the left the color on the inflated cortical surface of the left hemisphere shows the cortical thickness of the subject, obtained with micapipe, expressed as a z-score relative to a cohort of 160 healthy controls. The regions with no shading indicate where the activity recorded by the iEEG is generated. A negative correlation is observed with log-transformed spike rates (r=0.32), significant in a block permutation test preserving the local correlations of the cortical thickness (p=0.009). *Center*: There is no correlation between the source localization of an average spike (from 44 occurrences) at the location from which the iEEG channels record and the intracranial spike rate. The reason for the lack of correlation is that the scalp spikes represent neuronally propagated activity. The spikes originate in the mesiotemporal and mesial occipital regions and then propagate to the lateral temporal region, giving rise to the activity seen in the scalp EEG. The possible propagation path can be obtained from tractography. The center and bottom left show the tracts passing within 5 mm of the iEEG contacts of the hippocampal channel LH1-2 and the cortex (only left hemisphere shown); this allows obtaining the anatomical cortical connectivity map for each iEEG channel. Comparing these maps to the source imaging solution, we obtain a back-propagated activity for each channel (bottom center). This backpropagated scalp source activity recovers a positive correlation with the intracranial epileptic spike rate (r=0.3, p=0.035, block permutation test).

The middle-center panel of **Figure 4** shows electro-MICA source localization results for the average of 44 epileptic left temporal spikes on the scalp EEG, at the peak of the spike, for good SNR. The middle-right panel shows that no significant correlation is found between the average source localization strength at the regions from which the iEEG channels record and the log-spike rate (r=0.02; p=0.9, block permutation test preserving the local correlations, 10^5^ permutations). This is not surprising, since in this patient the scalp spikes are propagated activity from deeper generators in the hippocampus and mesial occipital region. The epileptic spikes from deep generators propagate to the neocortex following white matter tracts (Mitsuhashi et al., 2021; Azeem et al., 2023). From the anatomical connectome obtained with micapipe, we can obtain cortical maps showing the strength of the connections between the regions from which the iEEG channels record and the whole cortex. The lower left panels of **Figure 4** show an example of the main white matter tracts between the proximity of the contacts of intracranial channel LH1-2 and the rest of the cortex. This channel records from the hippocampus and has one of the highest epileptic spike rates. We can see that it connects to the lateral temporal lobe and to the occipital lobe. We can estimate the contribution from each iEEG location to the scalp spike through propagation by computing the spatial correlation between its cortical connectivity map and the source localization map, obtaining one value per iEEG channel, shown projected on the inflated cortex in the bottom-center panel of **Figure 4**. The bottom-right panel shows the significant positive Pearson correlation between this back-propagated scalp spike and the log-spike rate (r=0.30; p=0.035, block permutation test preserving the local correlations, 10^5^ permutations). In other words, the backpropagated scalp spike source is positively correlated with the intracranial spiking, as expected.

## 4. Discussion

We presented electro-MICA, a toolbox for integrating electrophysiology data from scalp or intracranial EEG with multimodal imaging and connectomics results. The toolbox is a companion to micapipe and HippUnfold, extending its scope as a multimodal imaging ecosystem. It is an open access toolbox purely in python and extremely user friendly, with no parameter selection needed. It is based on the numerical solution of physical models for the electric phenomena in the brain. We presented data supporting our choice of models, and we showed an example of the association between pathological electrophysiology and cortical thickness, and the ease of integration with connectivity results from micapipe. Other examples of possible uses would be the study of associations between microstructural features, connectivity, and intracranial (Royer et al., 2025) or scalp (Sahlas et al., 2024) epileptic spikes, or the association between fMRI derived hemodynamic features and high frequency oscillations (Zhou et al., 2024).

We prioritized ease of use for the pipeline, without compromising the quality of the results. This means that we did not include advanced source localization algorithms that might require more user intervention and tunning of parameters. The intention is to provide seamless integration with other imaging modalities, not another software package for EEG processing, as there are powerful existing solutions for this such as MNE (Gramfort, 2013), EEGlab (Delorme & Makeig, 2004), or brainstorm (Tadel et al., 2011). However, the iEEG sensitivity map, or leadfield, produced by electro-MICA incorporates unique models that make it particularly appropriate for iEEG source localization. In particular, iEEG source localization studies (Juárez-Martinez et al., 2018; Cimbalnik et al., 2019; Medina Villalon et al., 2024) could benefit from the non-singular model for the generator we included in this pipeline.

We encourage the community to explore electro-MICA and to contribute to future enhancements of this toolbox. Future versions may include more tissue classes in the model and allow anisotropy for the white matter and skull bone. The differentiation of cerebrospinal fluid (CSF), gray matter and white matter, while not essential, has been shown to improve iEEG modeling (Von Ellenrieder et al., 2012). Incorporating CSF and allowing for skull anisotropy would also improve the scalp EEG model (Wolters et al., 2006). Computation of the models could be accelerated by using the inskull surface, as well as scalp electrode positions if available, to predict the shape of other surfaces (Valdés-Hernández et al., 2009), since morphological operations in 3D are comparatively slow. Additionally, if the position of the scalp electrodes is provided, the BIDS requirements currently mandate a coregistered anatomical image. This requirement could be eased by coregistering the electrodes to the micapipe space directly.

Electro-MICA is a toolbox for the integration of electrophysiology and multimodal neuroimaging within a unified, accessible framework. Grounded in physical models, the toolbox achieves a balance between methodological robustness and practical simplicity. The validation results demonstrate the superiority of the distributed generator model over distance-based and dipole-based alternatives, and the clinical example illustrates how electro-MICA can reveal meaningful relationships between pathological electrophysiology, cortical structure, and brain connectivity that would otherwise require significant custom development to uncover. As an open-access, parameter-free Python toolbox fully compatible with the micapipe/HippUnfold software ecosystem, electro-MICA lowers the barrier for researchers and clinicians seeking to combine the temporal richness of EEG with the spatial precision of modern neuroimaging. We anticipate that electro-MICA will serve as a valuable platform for the multimodal investigation of both healthy brain function and neurological disorders.

## Data AND Code Availability

The toolbox is available at http://github.com/MICA-MNI/electromica with extensive online documentation at http://electromica.readthedocs.io.

## Author Contributions

NvE conceptualization, data curation, methodology, software, visualization, writing—original draft; ZC conceptualization, software, writing—reviewing & editing; TA data curation, writing—reviewing & editing; LV formal analysis, visualization, writing—reviewing & editing; CA data curation, writing— reviewing & editing; JDK software, writing—reviewing & editing; RRC software, writing—reviewing & editing; JR conceptualization, writing—reviewing & editing; ES conceptualization, writing—reviewing & editing; PB conceptualization, writing—reviewing & editing; RP resources, writing—reviewing & editing; OA resources, writing—reviewing & editing; BF resources, writing—reviewing & editing; BCB conceptualization, funding acquisition, project administration, resources, writing—reviewing & editing.

